# Resolving Tumour Clonal Heterogeneity and Spatial Complexity using Nuclear Tandem Epitope Protein (nTEP) Barcoding

**DOI:** 10.1101/456855

**Authors:** Michael B. Gill, Simon Koplev, Anne C. Machel, Martin L. Miller

## Abstract

Tumours are composed of an array of unique cancer cell clones along with many non-tumour cells such as immune cells, fibroblasts and endothelial cells, which make up the complex tumour microenvironment. To better understand the co-evolution of tumour clones and cells of the tumour microenvironment, we require tools to spatially resolve heterotypic cellular interactions at the single cell level. We present a novel protein-based barcoding technology termed nuclear tandem epitope protein (nTEP) barcoding, which can be designed to combinatorially encode and track dozens to hundreds of tumour clones in their spatial context within complex cellular mixtures using multiplexed antibody-based imaging. Here we provide proof-of-principle of nTEP barcoding and develop the technology, which relies on lentiviral - based stable expression of a nuclear-localised fluorophore that contains unique combinations of protein epitope tags that can be decoded by a limited set of antibodies. By generating a series of cell lines expressing unique nTEP barcodes, we were able to robustly identify and spatially deconvolve specific clones present within highly complex cell mixtures at the single cell level using state-of-the-art iterative indirect immunofluorescence imaging (4i). We define the utility of nTEP-barcoding as a powerful tool for visualising and resolving tumour heterogeneity at the cellular level, and envision its usage in mouse tumour models for understanding how tumour clones modulate and interact with stromal- and immune cells in cancer.

## INTRODUCTION

Tumours are complex and dynamic entities composed of multiple unique mutant sub-clones^1^ and a large array of non-tumour cells including immune and stromal cells.^2,3^ It is of significant interest to the field of cancer biology to understand the complex interplay between cells within the tumour microenvironment (TME) and how different cell populations spatially alter during tumour progression.^2–4^ At present, considerable efforts are underway to harness the power of multiplexed methods to examine the dynamics of the TME. For example, single-cell RNA sequencing can resolve individual cell types, measuring thousands of transcripts in tumours dissociated into single cell suspensions^5,6^; merFISH (multiplexed error-robust fluorescence *in situ* hybridization) can examine the expression of 100-1000 of transcripts within individual cells within complex tissue sections^7–9^; IMC (imaging mass cytometry) can identify dozens of specific proteins and their cell type-specific expression within tumour sections.^10^ Additionally, with the recent publication of the a new imaging platform, termed iterative indirect immunofluorescence imaging (4i)^11^, it is now possible to image the spatial distribution of at least 40 proteins using multiple rounds of stripping, re-staining, and immunofluorescence imaging. Although powerful, none of these technologies currently provides simultaneous tracking of individual tumour clones resolved in their spatial context of immune and stromal cells of the TME during tumour progression in mice.

In order to fully harness the potential of imaging methods such as 4i and IMC to track individual tumour clones in the spatial context of the TME at the single cell level, we propose a new protein barcoding system called nuclear tandem epitope protein-barcoding (nTEP-barcoding). This technology uses a lentiviral expression of a central nuclear localised fluorophore, e.g. H2BeGFP, which is tagged in combination with small linear epitopes selected from a larger library of epitopes. The tandem encoding of a fluorophore and a combinatorial set of epitopes creates a straightforward way of expanding the system by exchanging the core fluorophore (e.g. eGFP to mCherry), thereby doubling the set of possible barcodes. This constitutes a highly multiplexed barcoding system, which can be deconvoluted with a small number of antibodies, thus permitting the simultaneous detection of barcoded clones as well as proteins related to key aspects of cancer biology such as signal transduction and immune cells. Another key feature of the methodology is that nTEP barcoding uses a nuclear localised H2B-fusion protein as the core barcoding unit, enabling greater clarity for 4i or IMC analysis as barcoded cells are readily segmented (nucleus delinarised) and distinguished from non-cancer cells in complex tumour tissues.

Simultaneous with this paper, a method termed Pro-Code that utilises plasma membrane-based epitope barcodes was published.^12^ Pro-Code enables highly multiplexed protein-based barcoding of cell clones and was applied in conjunction with single-cell CRISPR screens using flow-based mass cytometry. Although suitable for analysis of clonal dynamics and signalling states at the single cell level in cellular suspensions, such methods are not useful for spatial compartmentalisation and imaging of barcoded clones within complex cellular mixtures or tissues due to close cell-cell membrane contacts resulting in overlapping, indistinguishable cell populations.

Here, we provide a proof-of-concept of a new protein-based barcoding system of cell clones. The barcodes can be spatially resolved in complex cell mixture at the single cell level using 4i imaging. Future studies will apply nTEP barcoding in mouse tumour models which will enable first-of-its-kind high-dimensional, single cell-based map of individual cancer clones and their proximity to stromal and immune cells during *in vivo* tumour evolution.

## RESULTS

### nTEP barcoding for studying clonal heterogeneity in the tumour microenvironment

We developed a protein-based barcoding system by using lentivirus-based stable expression of unique epitope-fluorophore fusion proteins for single-cell visualisation of barcoded tumour clones (**Figure 1**). We generated unique, nuclear-localised fusion protein barcodes by using combinations of linear epitope tags (e.g. Myc, HA, V5 etc.) expressed in tandem with a H2Btagged fluorophore (e.g. eGFP or mCherry). The combinatorics of nTEP barcoding follows *n*choose-*k*, where *n* is the number of linear epitopes in the library and *k* is the selected set of epitopes chosen (**Figure 1A**). In addition, the expressed fluorophore not only provides efficient identification of nuclei of barcoded cells but also the total number of barcode combinations, *C*, can be expanded through exchanging the central fluorophore (*s*) creating a new set of constructs through a straightforward cloning step (e.g. eGFP to mCherry, see **Figure 1A** for details). For *in vitro* experiments, nTEP can be decoded by a limited number of antibodies equal to the library of epitopes (*n*) as the fluorophore signal is retained in fixed of cell lines, while additional antibodies may be needed to detect the fluorophore in FFPE fixed tissue samples. The principle behind nTEP is similar to the recently published Pro-Code method^12^, but nTEP adds additional encoding through the exchangeable fluorophore as well as the advantage of *in situ* analysis in a tissue context such as the tumour microenvironment.

**Figure 1.**
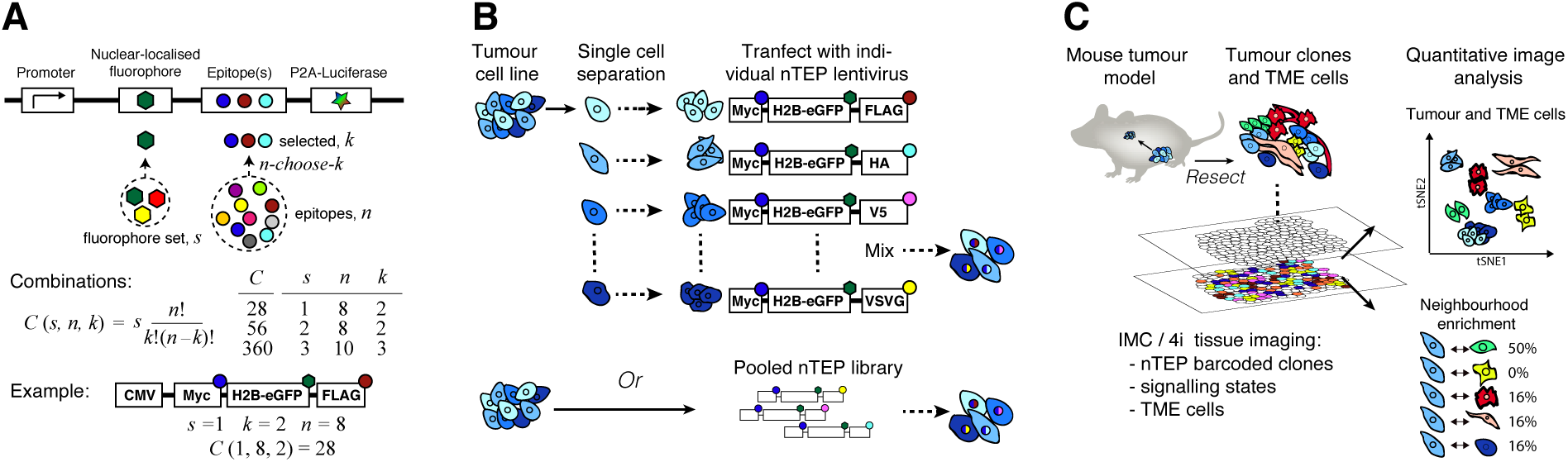
Design and utility of nuclear Tandem Epitope Protein (nTEP)-barcoding. (**A**) Schematic illustration of the nTEP expression cassettes and barcoding principle comprising tandem expression of a nuclear localised fluorophore with combinations of epitope tags. A number of epitopes (*k*) are selected from a library of epitopes (*n*) in an n-choose-k fashion. The total number of combinations (*C*) is further expanded by using additional core fluorophores (*s*) creating a new sets of constructs. For example, using two epitopes selected from a library of eight epitopes and one fluorophore would provide 28 distinguishable tags [C(*s*, *n*, *k*) = C(1, 8, 2) = 28]. This can be expanded with another set of 28 to 56 using an additional fluorophore [C(2, 8, 2) = 56]. (**B**) Lentiviral-based stable expression of nTEP barcodes using an individual or pooled approach. (**C**) Application of nTEP for single cell analysis of injected barcoded cell lines using *in vivo* mouse tumour models and multiplexed antibody imaging such as IMC or 4i. In addition to the barcoded clones themselves, key features of tumours such as signalling pathways and TME cells can be analysed through antibody measurements to ultimately investigate spatial complexities of co-evolving tumour clones and TME cells.

The nTEP barcoding method can be used in a pooled manner through low-titer infection of an nTEP lentiviral library or by transfecting individual lentiviral barcodes to cell lines as exemplified by cells lines derived from a heterogeneous tumour cell line (**Figure 1B**). In the current version, we focussed on developing and benchmarking nTEP using barcoding of individual cell lines with unique barcodes and by using combinations of single and dual epitopes added to the N- or C-terminal end of a H2B-eGFP cassette. Each unique nTEP-barcode was cloned into a lentivirus vector and used to generate viral stocks. The resulting viruses were used to infect cells and generate stable cell lines, which could be analysed for the expression of the single epitope tags as well as for the detection of all epitopes in more complex cell mixtures. We envision developing nTEP barcoding for tracking how genomically or phenotypically heterogeneous tumour clones behave in syngeneic tumour mouse models by mixing the barcoded cells and injecting them subcutaneously or orthotopically into mice (**Figure 1C**). By using state-of-the-art imaging with multiplexed antibody measurements, such as IMC or 4i, the spatial complexity of barcoded cellular mixtures can be deconvoluted with a limited set of antibodies leaving multiple antibodies to measure signalling processes and stromal- and immune cells of the TME.

### Selection and verification of epitopes for nTEP barcoding

A panel of published epitope tags were selected based on both the relative size of the antigenic sequence, with shorter tags being favoured, and their published use in the literature. Initially, we selected and tested 10 epitope tags including FLAG, Myc, V5, VSVG, HA, HIS, AU1, B-tag, Glu-Glu and HSV1. Primers were generated in order to amplify a H2B-eGFP encoding cassette, which encodes a nuclear localised fluorescent protein, placing an epitope tag at the 5’-primer region of the H2B-eGFP, generating (X)-H2B-eGFP, where X is the epitope tag. The amplified products were cloned into the lentivirus expression plasmid pLVX-Puro using incorporated restriction sites. The resulting fusion proteins were expressed in HEK293T cells and stained for the expression of their epitope tag using immunofluorescence microscopy (**Figure 2A**). All of the epitope-tagged fusion proteins exhibited clear expression and nuclear localisation, demonstrating that the presence of the 5′-primer tag did not alter the stability or nuclear localisation of H2B-eGFP. Although the HIS tag in the HIS-H2B-eGFP fusion protein was mainly localised in the nucleus (data not shown) it also demonstrates a high level of non-specific cytoplasmic staining, thus HIS was omitted in this and from future studies.

**Figure 2.**
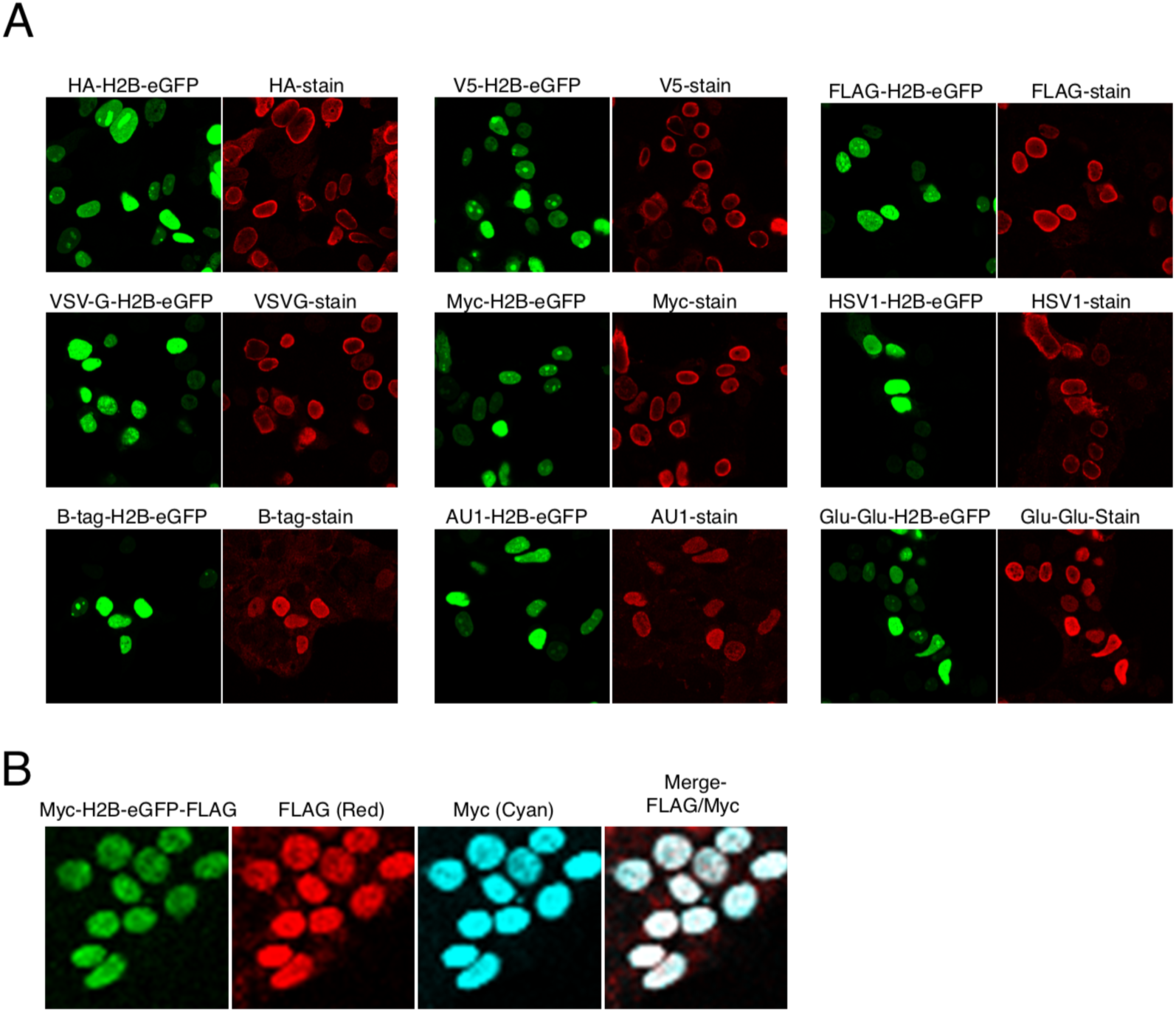
Immunofluorescence staining shows nuclear localisation of expressed nTEP-barcodes. Representative images of stable HEK293T cells expressing either single (**A**) or double (**B**) tagged nTEP-barcodes stained for the presence and nuclear localisation of the respective epitope tag. The detected epitope co-localises with H2B-eGFP and does not alter the nuclear targeting of the expressed fusion protein.

We next examined the impact of dual tagging the H2B-eGFP cassette to ensure that both tags could be detected simultaneously without compromising the nuclear targeting of the fusion protein. As an example, HEK293T cells expressing a 5′ Myc tag and 3′ FLAG tag were generated and used for immunofluorescent staining for both Myc and FLAG (**Figure 2B**). The presence of the dual tag did not alter nuclear targeting and both tags would be co-stained.

We further tested if the selected panel of antibodies demonstrated cross reactivity given that the onward experiments would utilise a combination of cells expressing either a single or double tagged-H2B-eGFP barcodes. To do this a panel of six cell lines were selected, each expressing a unique tag (HA, FLAG, Myc, V5, VSVG or HSV1). Each cell line was stained independently using all six antibodies analysed using immunofluorescence microscopy. No apparent cross-reactivity was observed (**Figure S1**).

### Robust identification of singly-barcoded cells within mixtures of cell lines

We next sought to show the detection of single barcodes within a mixture of cells. Six independent cell lines expressing a single barcode (FLAG, HA, VSVG, Myc or V5), and a cell line expressing only H2B-eGFP, were equally mixed, plated and grown for two passages. To detect the presence of three of the total nTEP-barcoded cells present in the mixture, cells were first stained for HA, FLAG and VSVG (**Figure 3A**). Merging images across the three stainings using Fiji imaging software indicated that the single nTEP-barcodes could be clearly distinguished as separate cell populations. As expected, there was limited overlap between the barcodes of the individually cells.

**Figure 3.**
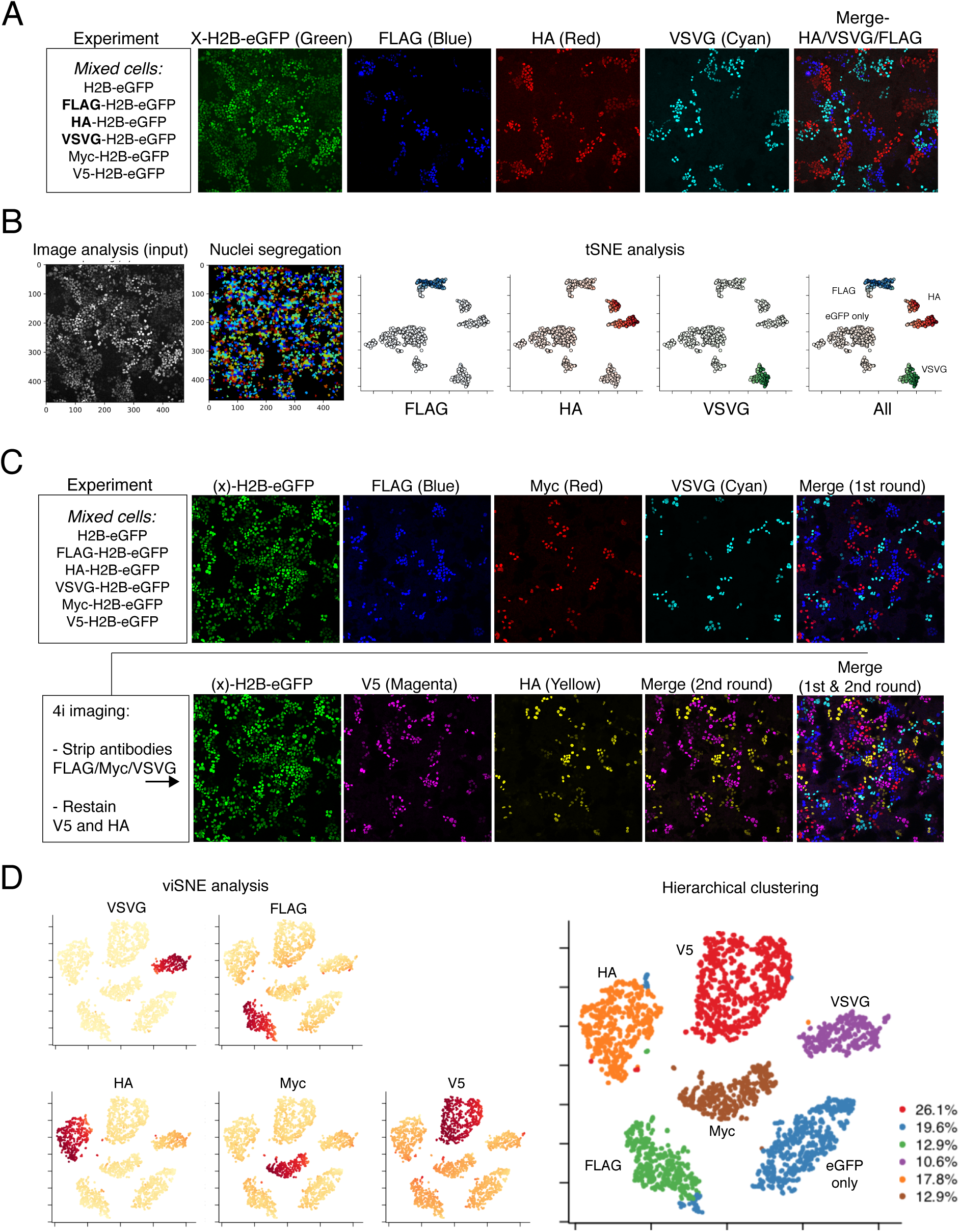
Single epitope nTEP-barcoded cell lines are distinctly identified in complex cellular mixtures. (**A**) Mixed cell populations containing six independent single tagged nTEP-barcodes (including a H2B-eGFP non-tagged control) were imaged and analysed to define the spatial positioning by staining for three barcodes (FLAG, HA and VSVG). (**B**) Quantitative image analysis was applied to define and segreate nuclei using H2B-eGFP and further analysed for barcode representation using tSNE. Only cells within the 50th percentile H2B-eGFP intensity were included, for a total of n=826 cells, and coloured proportional to the mean intensity antibody stain for each epitope. Colours were mixed by their minimum RGB values. (**C**) In order to analyse all six barcoded cell lines in the spatial context, 4i was employed with one round of elution and restaining the slides enabling imaging of all nTEP barcodes. (**D**) Pooled barcode analysis from three multiplexed microscopy views covering n=3,271 single cells. The composition of the individual clone sets was analysed by viSNE. Hierarchical clustering was performed yielding six distinct populations corresponding to the six clones mixed.

To quantitatively analyse the mixtures of barcoded cells, we performed single-cell image analysis using CellProfiler.^13^ We first defined nuclei using the H2B-eGFP image, segregating individual cells and subsequently estimated the mean nuclei intensities of the epitope stains. To assess the structure of the multiplex epitope stains, we performed tSNE dimensionality reduction of epitope intensities for cells with high H2B-eGFP expression (n=826) (**Figure 3B**). These data show that singly-barcoded cell lines (FLAG, HA and VSVG) separate both from each other by their unique nTEP barcode and from the bulk population of cells, which contains a mixture of eGFP only and barcoded cells not directly measured (Myc and V5).

To enable the detection of all nTEP barcodes expressed by individual cells within the mixed cell population, we employed the elegant 4i strategy for iterative antibody staining of the same slide. Initially, the mixture of cells were stained to detect spatial position of individual cells expressing FLAG, Myc and VSVG (**Figure 3C**, upper panel). The initial staining antibodies were eluted using the 4i protocol and the cells were re-stained to detect the spatial localisation of cells expressing V5 and HA (**Figure 3C**, lower panels). In order to superimpose the spatial positioning of all singly-barcoded cell populations the images for each independent antibody stain were pseudo-coloured and merged using Fiji imaging software (**Figure 3C**). The spatial distribution of the single nTEP-barcoded cell populations were analysed by registrering (aligning) H2B-eGFP images across 4i rounds, performing nuclei segmentation, and quantifying multiplex epitope stains. The mean epitope intensities were pooled across three microscopy views and analysed by viSNE, which yielded six well-defined populations including the H2BeGFP only population, as expected from the experimental setup (**Figure 3D**). Hierarchical clustering further defined six cell populations with comparable fraction of barcoded cells except V5, which had higher intensity stains likely resulting in high recovery rates in addition to stochastic events such as variable growth rates.

### Combinatorial and single nTEP-barcoded cell populations decoded from cell mixtures

To further extend our analysis we also examined an equal mixture of nine cell lines tagged with six single nTEP barcodes (FLAG, HA, VSVG, V5, Myc or HSV1) and three combinatorially encoded cells (Myc-HA, Myc-VSVG and VSVG-HA). The plated cells were fixed and stained for the presence of all six epitopes using 4i with initial staining for FLAG, HA and VSVG, followed by antibody stripping and restaining for V5, and a last round for Myc and HSV1 (**Figure 4A**). Merged images for all six epitope tags revealed both presence of single and double positively stained nuclei as expected. To decode the single and combinatorially barcoded cells, we applied image analysis using a similar approach as described for the singly-barcoded cells. Overall the nine barcoded populations had excellent separation (**Figure 4B**) taking into account a false positive rate of approximately 1% (**Supplementary Figure S2**).

**Figure 4.**
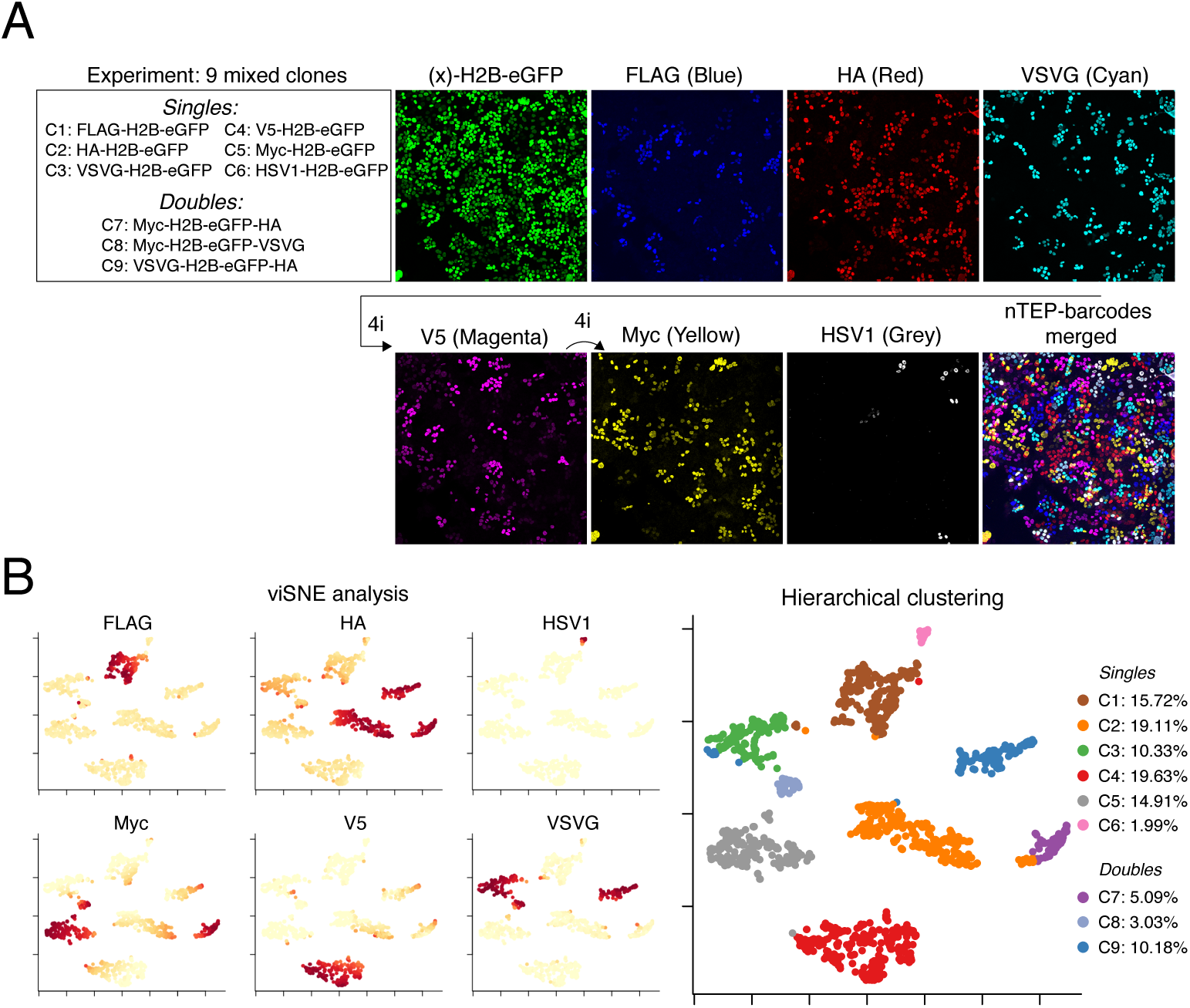
Identification of single and combinatorially tagged nTEP-barcodes in complex cellular mixtures. (**A**) Mixed cell populations containing six independent single tagged nTEP-barcodes and three dual tagged nTEP-barcodes were imaged and analysed to define the spatial positioning by staining for the six epitopes expressed using two rounds of 4i (FLAG, HA, VSVG, V5, Myc and HSV1). (**B**) The composition of the individual clone sets was analysed by viSNE and hierarchical clustering was performed yielding nine distinct populations corresponding to the nine clones mixed. A total of n=1,355 cells were analysed.

## DISCUSSION

The heterogeneous nature of tumours not only poses great challenges for understanding and treating cancer but also for creating model systems that can capture the complexity of cancer at the molecular, phenotypic and cellular levels. To understand the complexity of the TME various technologies have been developed for labelling and tracking individual cells present within complex mixtures. Elegant studies have used DNA barcoding technologies to evaluate the presence of unique tumour clones present within the primary tumour and present at distal metastasis.^4^ Although sophisticated, these studies do not enable the spatial complexity of the unique clones to be defined or how heterogeneous tumour clones modulate TME cells. Analysis of single cells using both RNA sequencing and protein barcoding with Pro-Code (using CyTOF) also enables detailed analysis of individual clones but crucially lacks spatial context of investigated cells.^5–7,12^

With advances in protein-based imaging platforms, including IMC and 4i, it is now to possible to analyse dozens of proteins within complex cellular mixtures or tumour sections. Although powerful, these technologies have so far mainly been applied to generically detect proteins of interest and are rarely used to distinguish between closely related cells such as different tumour clones originating from cancer stem cells. To harness the power of these new technologies, we have here developed a new protein barcoding technology that enables spatial tracking clones present within complex cellular mixtures using basic antibody-based imaging strategies.

In this study, we have demonstrated that individual cells labelled using specific nTEP barcodes can be identified as unique nuclei when present in complex cellular mixtures. The nuclear localisation of nTEP provides excellent cell segmentation for defining individual cells when present in more closely packed cellular structures. This is illustrated by the fact that we find low error rates (<1%) when estimating the double positive error rates among mixtures of three single-epitope nTEP-barcoded cells *in vitro* (**Supplementary Figure S2D**), although error rates are likely to increase in more complex *in vivo* tissues. The use of fluorophores in tandem with the epitope encoding will also allow for future use of fluorescence-guided IMC to narrow the search space for IMC, which can otherwise be limited by long imaging requirements. Furthermore, utilising additional fluorophores constitutes a clear approach to expanding the barcoding system by exchanging the core fluorophore and doubling the set of possible barcodes. Promisingly, the use of 4i provides a new way of multiplexed antibody-based imaging, and we find very low spillover rates (<0.2%) from consecutive stain and restain processes (**Supplementary Figure S2C**).

In contrast to most DNA and RNA single cell-based technologies, which require dissociation of tumour tissues in to cell suspensions, the nTEP barcoding technology enables cells to be examined in their physiological context. For example, by implanting a unique set of nTEP-barcoded tumour cell clones, using established tumour cell based models, it will likely be possible to identify single-cell cancer clones within its tumour microenvironment using only a limited set of antibodies. When used in conjugation with 4i and IMC, investigators will not only be able to identify the relative cellular positioning and competition of the implanted barcoded tumour cells, but also pinpoint the relative positioning of the non-tumour clones within the surrounding microenvironment (e.g. immune cells and supporting stromal cells). Using such imaging technologies in conjugation with drug perturbations studies would lead to a greater understanding of how therapeutic drugs act upon or even change the spatial composition of both the individual tumour subclones and non-tumour cells present within the TME.

## METHODS

### Cells

HEK293T were all maintained at 37 °C with 5% CO_2_. All cells were grown in DMEM-F12 with 10% FBS. Stable HEK293T cell lines expressing nTEP barcodes were generated by infecting cells with individual nTEP barcode lentiviruses and selection with 3 μg/ml Puromycin.

### nTEP barcodes

H2B-eGFP was amplified using primer sets which incorporate as 5′ or 3′ epitope tag sequence. Amplified barcodes were cloned into the lentivirus clone pLVX-Puro using 5′ and 3′ incorporated restriction sites. Each clone was sequenced to ensure correct incorporate of the 5′, 3′ or both 5′ and 3′ epitope tags.

### Virus generation

Each pLVX-Puro-nTEP barcode clone was transfected into HEK293T cells along with the third generation packaging vectors (pMDL, pCMV-Rev and pVSV-G) using Lipofectamine 2000 (Thermo Fisher Scientific). Virus was collected, filtered (0.45 μM pore) and stored at −80 °C.

### Cell staining (immunofluorescence)

Single cells or mixed cell populations were either plated in poly-Lys treated cover glass or grown in chamber slides (u-Slide 2 well, #1.5H (170 μm plus-minus 5 μm)) (Integrated BioDiagnostics, Catalogue number 80287). Once grown to 90% confluency the cells were fixed using 4% formaldehyde for 20 m at room temperature (RT). Post fixation the cells were permeabilised using 0.1% Triton-X100/PBS for 15 m at RT, washed with 1X PBS and then blocked using 3% BSA/PBS/0.1% Tween (Blocker) for 1 h at RT. All primary antibodies were used at 1 in 500 dilution except [Y69] to c-Myc (Alexa Fluor^®^ 647) and FLAG-(29E4.G7) [DyLight 405] which were used at 1 in 100. Primary antibodies were incubated in blocker for 1 h at RT and then extensively washed with 1X PBS/0.1%Tween (TPBS). Secondary antibodies were all used at 1 in 1000 dilution in blocker for 1 h at RT and then washed using TBPS. Cover slides were mounted using ProLong™ Gold Antifade Mountant (Thermo Fisher Scientific, catalogue number P36930). Chamber slides were imaged directly without cell mounting.

### Iterative indirect immunofluorescence imaging (4i)

Cells immunofluorescently stained in chamber slides were imaged using confocal microscopy. In order to re-image the same cellular mixtures using a different set of primary antibodies the recently published 4i protocol was utilised to remove the first round of bound antibodies and enable re-staining.^11^

### Confocal microscopy and imaging processing

All images were captured using a Leica SP5 confocal microscope. Fiji software was used for image processing. All images were saved in a lossless format for further analysis.

### Image registration and nuclei quantification

For the 4i experiments, to adjust for small changes to rotation and XY-coordinates after consecutive acquisition of confocal microscopy images, we used a Discrete Fourier Transform (DFT) method^14^ to align the H2B-eGFP channels. We used the Python package imreg_dft 2.0.0 with the optional argument set at 10 iterations and with the default pixel-by-pixel alignment. The H2B-eGFP images were converted to grayscale before alignment and the estimated parameters for XY offset, scale, and rotation were used to translate all antibody and eGFP stains per iteration.

Next, we used CellProfiler 3.1.5^13^ to define cell nuclei based on the H2B-eGFP channel. We used the global threshold strategy for minimum cross entropy without threshold smoothing. Detected nuclei with diameters less than 5 or larger than 12 pixels were excluded, as were nuclei bordering the edge of the image. For the remaining nuclei, we then calculated the mean intensity of each antibody and H2B-eGFP channel.

### Multiplex nuclei clustering

Unless otherwise stated we log transformed the mean nuclei intensities, clamping values below 0.01. tSNE analysis was performed with the tsne R package and viSNE^15^ with the Rtsne package, which is a wrapper for a C++ implementation of the Barnes-Hut algorithm.^16^ Both methods were run for 1,000 iterations. Hierarchical clustering was performed with complete linkage and the Euclidean distance metric. To estimate cell type frequencies the tree was cut at the expected number of clusters and each cluster was manually annotated with corresponding barcodes.

Primary antibodies used in this study.

Primary antibodies used in this study
| Primary Antibody to<br>(or conjugated) | Host species | Supplier | Catalogue number |
| --- | --- | --- | --- |
| [Y69] to c-Myc (Alexa Fluor® 647) | Rabbit | Abcam | ab190560 |
| Myc (Biotin) | Rabbit | Abcam | ab34773 |
| Myc | Mouse | Millipore | 05-724 |
| Flag (clone M2) | Mouse | Sigma | F1804 |
| FLAG-(29E4.G7)<br>[DyLight 405] | Mouse | Bio-Techne | NBP1-97399 |
| HA | Mouse | Abcam | ab130275 |
| HA | Rabbit | Sigma | H6908 |
| VSVG | Mouse | Abcam | ab50549 |
| VSVG | Rabbit | Abcam | ab1874 |
| V5 | Mouse | Abcam | ab27671 |
| HSV1 | Rabbit | Genscript | A00624-40 |
| AU1 | Rabbit | Genscript | A00628-40 |
| B-tag | Rabbit | Genscript | A00638-40 |
| Glu-Glu | Rabbit | Genscript | A00627-40 |

Secondary antibodies used in this study.

Secondary antibodies used in this study
| Secondary Antibody | Supplier | Catalogue number |
| --- | --- | --- |
| Donkey anti-Rabbit IgG (H+L)<br>Highly Cross-Adsorbed<br>Secondary Antibody, Alexa | ThermoFisher SCIENTIFIC | A10042 |
| Fluor 568 |  |  |
| Goat anti-Rabbit IgG (H+L)<br>Highly Cross-Adsorbed<br>Secondary Antibody, Alexa<br>Fluor 633 | ThermoFisher SCIENTIFIC | A21071 |
| Goat anti-Mouse IgG (H+L)<br>Cross-Adsorbed Secondary<br>Antibody, Alexa Fluor 568 | ThermoFisher SCIENTIFIC | A11004 |
| Goat anti-Mouse IgG (H+L)<br>Cross-Adsorbed Secondary<br>Antibody, Alexa Fluor 633 | ThermoFisher SCIENTIFIC | A21050 |
| Streptavidin, Alexa Fluor™<br>555 Conjugate | ThermoFisher SCIENTIFIC | S-32355 |

## ACKNOWLEDGEMENTS

This research was supported by Cancer Research UK core grant (C14303/A17197) as well as Cancer Research UK pioneer award (RG97051).

## SUPPLEMENTARY FIGURES

**Figure S1.**
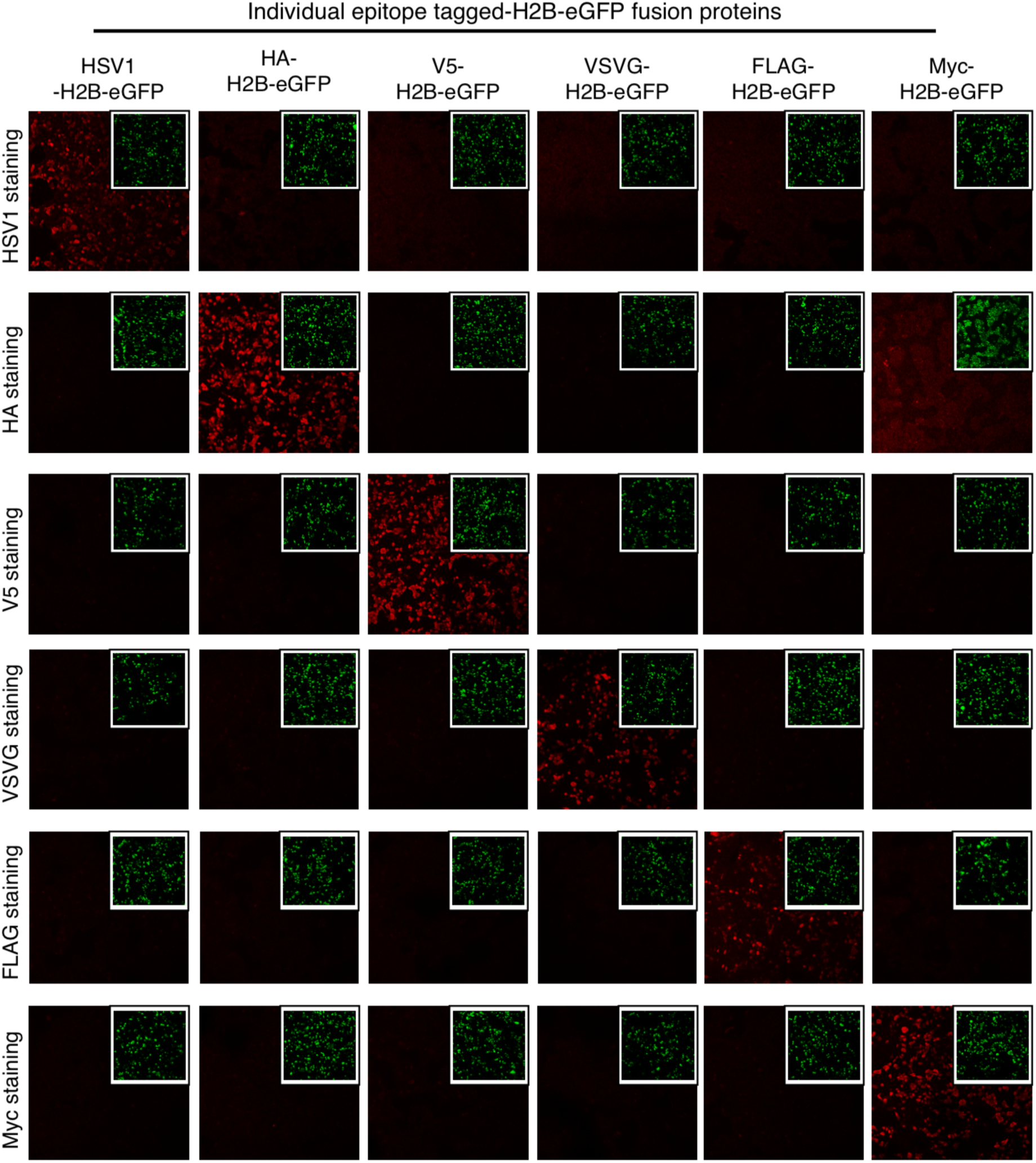
No cross-reactivity apparent using single epitope tagged nTEP-barcodes. HEK293T cells expressing either single epitope tagged nTEP-barcodes stained in an all by all fashion using antibodies specific to the epitope tags.

**Figure S2.**
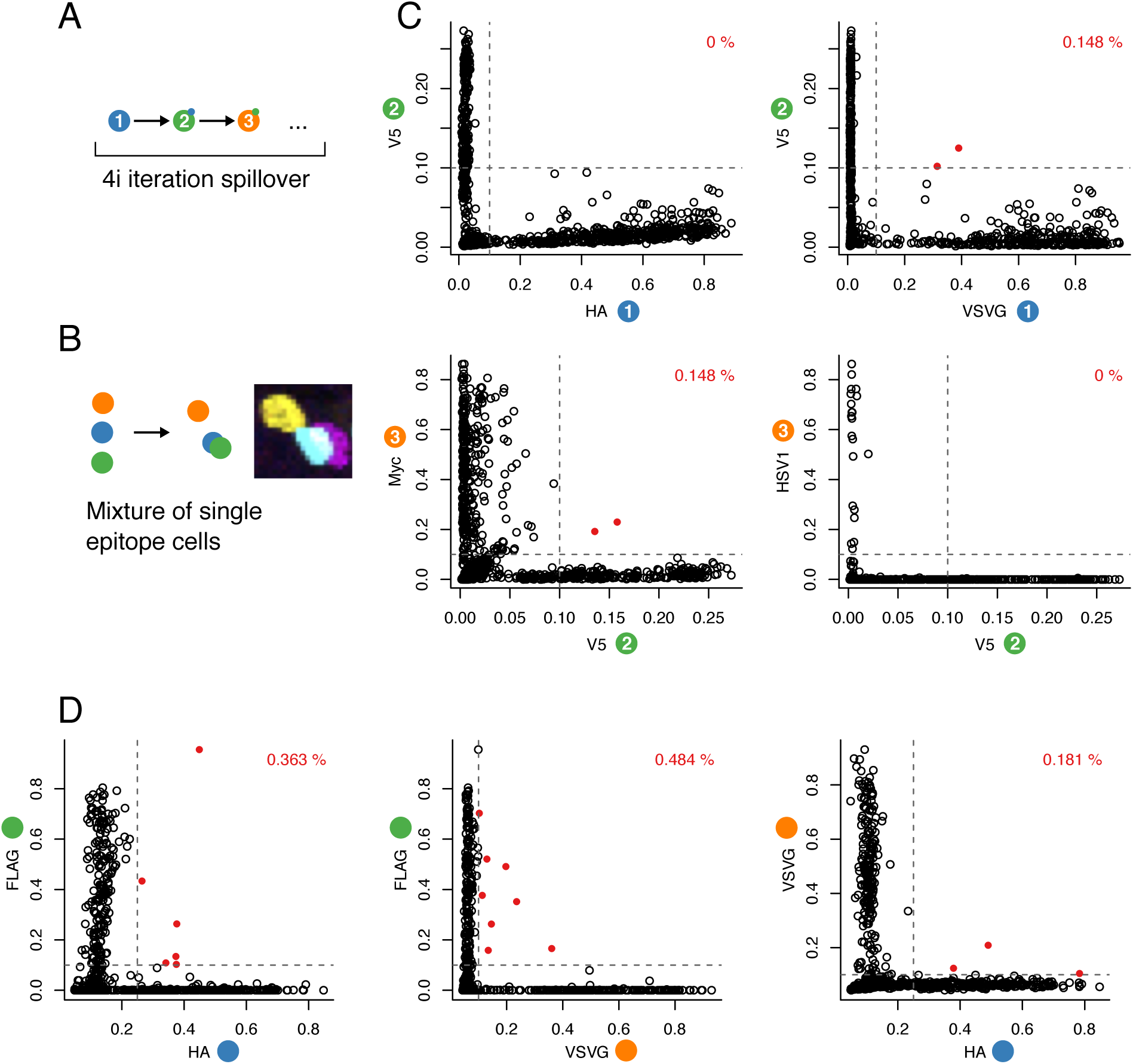
Estimation of false double positive rates of nTEP-barcoded cells due to spillover effects between 4i iterations and overlapping cells in culture. (A) Conceptual diagram of spillover stains in 4i iterations. (B) Example of how overlapping cells can lead to false double positive cells when nuclei definition is based on H2B-eGFP only. (C) Test of spillover stains in three iterations relating to Figure 4 in the main text. (D) Bottom three plots; estimates of double positive error rates among mixture of three single-epitope nTEP-barcoded cells relating to Figure 3 in the main text.

## REFERENCES

1. McGranahan, N. & Swanton, C. Clonal Heterogeneity and Tumor Evolution: Past, Present, and the Future. Cell 168, 613–628 (2017).

2. Quail, D. F. & Joyce, J. A. Microenvironmental regulation of tumor progression and metastasis. Nat. Med. 19, 1423–1437 (2013).

3. Joyce, J. A. & Fearon, D. T. T cell exclusion, immune privilege, and the tumor microenvironment. Science 348, 74–80 (2015).

4. Wagenblast, E. et al. A model of breast cancer heterogeneity reveals vascular mimicry as a driver of metastasis. Nature 520, 358–362 (2015).

5. Chung, W. et al. Single-cell RNA-seq enables comprehensive tumour and immune cell profiling in primary breast cancer. Nat. Commun. 8, 15081 (2017).

6. Levitin, H. M., Yuan, J. & Sims, P. A. Single-Cell Transcriptomic Analysis of Tumor Heterogeneity. Trends Cancer Res. 4, 264–268 (2018).

7. Chen, K. H., Boettiger, A. N., Moffitt, J. R., Wang, S. & Zhuang, X. RNA imaging. Spatially resolved, highly multiplexed RNA profiling in single cells. Science 348, aaa6090 (2015).

8. Moffitt, J. R. et al. High-throughput single-cell gene-expression profiling with multiplexed error-robust fluorescence in situ hybridization. Proc. Natl. Acad. Sci. U. S. A. 113, 11046–11051 (2016).

9. Moffitt, J. R. et al. High-performance multiplexed fluorescence in situ hybridization in culture and tissue with matrix imprinting and clearing. Proc. Natl. Acad. Sci. U. S. A. 113, 14456–14461 (2016).

10. Giesen, C. et al. Highly multiplexed imaging of tumor tissues with subcellular resolution by mass cytometry. Nat. Methods 11, 417–422 (2014).

11. Gut, G., Herrmann, M. D. & Pelkmans, L. Multiplexed protein maps link subcellular organization to cellular states. Science 361, (2018).

12. Wroblewska, A. et al. Protein Barcodes Enable High-Dimensional Single-Cell CRISPR Screens. Cell (2018). doi:10.1016/j.cell.2018.09.022

13. Carpenter, A. E. et al. CellProfiler: image analysis software for identifying and quantifying cell phenotypes. Genome Biol. 7, R100 (2006).

14. Reddy, B. S. & Chatterji, B. N. An FFT-based technique for translation, rotation, and scale-invariant image registration. IEEE Trans. Image Process. 5, 1266–1271 (1996).

15. Amir, E.-A. D. et al. viSNE enables visualization of high dimensional single-cell data and reveals phenotypic heterogeneity of leukemia. Nat. Biotechnol. 31, 545–552 (2013).

16. van der Maaten, L. Barnes-Hut-SNE. arXiv [cs.LG] (2013).

